# Focal exocytosis of Syntaxin 3 and TRPML1 at pseudopodia of nascent phagosomes

**DOI:** 10.1101/2022.08.22.504747

**Authors:** Deepti Dabral, Geert van Bogaart

## Abstract

Macrophages clear invading pathogens by phagocytosis. Phagocytosis is a complex mechanism involving the local expansion of the membrane, cytoskeletal remodeling, and the delivery of phagosomal proteins to the nascent phagosomes. However, the organelle trafficking events underlying this are largely unclear. Here, we show in human blood monocyte-derived macrophages that TRPML1, a calcium channel involved in the phagocytic process, is delivered to phagosomes in Syntaxin 3-positive vesicles. Syntaxin 3 is a SNARE protein previously shown to mediate the secretion of IL-6 by macrophages. Total Internal Reflection Microscopy (TIRF) revealed that Syntaxin 3 positive compartments carry TRPML1 to pseudopodia for focal exocytosis at the nascent phagosomes during *E. coli* uptake. Using siRNA knockdown, we show that both Syntaxin 3 and TRPML1 are required for *E. coli* uptake. Moreover, using TRPML1 agonists we show that increased TRPML1 activity leads to increased *E. coli* uptake, whereas calcium chelation decreased intracellular *E. coli* load. Understanding the membrane trafficking pathways is critical for understanding how macrophages clear invading pathogens.

**Key findings:**

1. Syntaxin 3 positive vesicles are delivered at the plasma membrane site of phagocytosis.
2. Syntaxin 3-positive vesicles carry TRPML1 to pseudopodia.
3. Bacterial phagocytosis correlates with Syntaxin 3 and TRPML1 expression levels.
4. Bacterial phagocytosis depends on calcium flux through TRPML1.
5. Syntaxin-3 vesicles carry the cytokine interleukin-6.

## Introduction

Macrophages clear invading pathogens and apoptotic bodies by phagocytosis. Phagocytosis is initiated by the binding of receptors on the macrophage plasma membrane to the phagocytic cargo particle. This activation of receptors triggers major cytoskeletal rearrangements to engulf the phagocytic cargo. The particle engulfment is accompanied by the focal exocytosis of endosomes and lysosomes to form plasmalemmal pseudopods. This focal exocytosis provides additional membrane and brings endo/lysosomal cargo molecules to the nascent phagosome (Bajno *et al*., 2000; Samie *et al*., 2013; Vashi *et al*., 2017; Sun *et al*., 2020). Already before the closure of the membrane, the newly-formed phagosomes undergo a maturation process that involves fusion with late endosomes and lysosomes, accumulation of lytic enzymes, acidification, and the generation of reactive oxygen species (ROS) by NADPH oxidases.

It is increasingly clear that many stages of phagocytosis are dependent on calcium signaling at the nascent phagosome (Nunes & Demaurex, 2010). For example, calcium promotes the activity of the NADPH oxidase NOX2, which mediates the killing of ingested microbes and antigen presentation (Bréchard *et al*., 2013). Moreover, calcium promotes the fusion of lysosomes with the plasma membrane (Czibener *et al*., 2006; Samie *et al*., 2013), where Synaptotagmin VII functions as a calcium sensor (Czibener *et al*., 2006). To phagocytose large particles (>3 μm), lysosomes undergo focal exocytosis in a Ca^2+^ dependent process (Medina *et al*., 2011; Samie *et al*., 2013; Xu & Ren, 2015), thereby providing extra membrane needed for the formation and extension of pseudopodia. Moreover, the release of lysosomal enzymes could also weaken the invading bacteria (Samie *et al*., 2013; Xu & Ren, 2015).

TRPML1 (Transient Receptor Potential Mucolipin-1) is an important calcium channel that promotes the phagocytosis of pathogens by focal exocytosis of lysosomes (Samie *et al*., 2013). The focal exocytosis of lysosomes at the forming phagosomes requires TRPML1 channel activity for supplying Ca^2+^ from its lumen (Samie *et al*., 2013). This focal exocytosis of lysosomes also requires the Ca^2+^-activated potassium (BK) channel for maintaining a membrane potential since an influx of K^+^ ions by the BK channel counters the out flux of Ca^2+^ ions by TRPML1 (Sun *et al*., 2020). The focal exocytosis of TRPML1 is mediated by VAMP7, a member of the soluble N-ethylmaleimide-sensitive factor attachment protein receptor (SNARE) protein family (Samie *et al*., 2013). However, the complete mechanism of how TRPML1 is focally exocytosed at nascent phagosomes is still incompletely understood.

SNARE proteins regulate exocytosis by forming complexes between cognate SNARE proteins in the plasma membrane and vesicular membrane (Dingjan *et al*., 2018). One of the cognate SNAREs of VAMP7 is Syntaxin 3 (Dingjan *et al*., 2018). Upon pathogenic activation of macrophages, Syntaxin 3 locates at the plasma membrane, where it interacts with the SNARE VAMP3 for secretion of the proinflammatory cytokine interleukin (IL)-6 (Frank *et al*., 2011; Collins *et al*., 2015; Dingjan *et al*., 2018). However, Syntaxin 3 is also located in intracellular compartments where it is involved in endosomal trafficking by interactions with VAMP8 (Dingjan *et al*., 2018). In resting macrophages, most Syntaxin 3 resides in intracellular compartments (Collins *et al*., 2015). Upon activation of the macrophages with the pathogenic stimulus lipopolysaccharide (LPS), Syntaxin 3 translocates to the plasma membrane likely for increased exocytosis of IL-6 and other factors (Collins *et al*., 2015). However, as LPS is mainly associated with bacteria, this raises the question of whether it would specifically translocate to phagocytic cups of nascent phagosomes and mediate focal exocytosis. Moreover, SNARE-mediated focal exocytosis of various organelles with the forming phagosomes to facilitate pathogen uptake has not only been observed for VAMP7-positive lysosomes (Samie *et al*., 2013), but also for VAMP3-positive recycling endosomes (Bajno *et al*., 2000) and Golgi-derived vesicles by VAMP2 (Vashi *et al*., 2017). As Syntaxin 3 interacts with VAMP2, VAMP3, and VAMP7 (Dingjan *et al*., 2018), we speculated that it could be involved in the focal exocytosis of TRPML1.

We, therefore, investigated whether Syntaxin 3 would mediate focal exocytosis to the nascent phagosomes for focal delivery of TRPML1 and uptake of *Escherichia coli* in macrophages. We show that Syntaxin 3-positive vesicles carrying TRPML1 are indeed focally delivered at the cups of nascent phagosomes in human peripheral blood monocyte-derived macrophages. This is essential for bacterial uptake, as siRNA silencing of either of these proteins reduces phagocytic uptake. In line with this, treating the macrophages with a TRPML1 agonist or an intracellular calcium chelator resulted in increased or decreased uptake of bacteria, respectively. Thus, the delivery of vesicles carrying Syntaxin 3 and TRPML1 is essential for the phagocytosis of *E. coli* by macrophages.

## Materials and Methods

### Mammalian cells and bacteria culture

Approval to conduct experiments with human blood samples was obtained from the Dutch blood bank Sanquin (Amsterdam, Netherlands) and all experiments were conducted according to national and institutional guidelines. Informed consent was obtained from all blood donors by the blood bank. Samples were anonymized and none of the investigators could ascertain the identity of the donors. CD14-positive monocytes were derived from buffy coats of healthy donors using the standardized Ficoll method [41]. Briefly, PBS containing 0.5 mM EGTA was added to the buffy coat, followed by centrifugation to collect the peripheral blood mononuclear cells. After extensive washing in 0.5 mM EGTA and 10% BSA, CD14-positive monocytes were collected using CD14-positive microbeads (Miltenyi Biotech, cat.no: 130-097-052) and magnetic antibody cell sorting (MACS) according to the manufacturer’s protocol. Monocytes were seeded at the density of 6 × 10^6^ cells in low attachment plates (Corning Costar, cat.no: 3471) and allowed to differentiate at 37°C, 5% CO_2_ for 8 days with 100 ng/ml macrophage colony-stimulating factor (M-CSF; R&D systems, cat.no 216-MC). Fully differentiated macrophages were harvested and seeded at the required density for the subsequent experiments.

*E. coli* strains XL 1 blue and K-12 MG1655 were cultured overnight in Luria Bertani (LB) broth media (Sigma Aldrich, cat. no: L3022). Overnight cultures were diluted to OD 600 ∼0.5 in RPMI1640 media supplemented with 10% fetal bovine serum. All stimulations were carried out at the ratio of 10 macrophages per bacterium, except for the colony forming unit (CFU) assays where macrophages were fed at a ratio of 50 macrophages per bacterium.

### Immunofluorescence Microscopy

15,000 macrophages were cultured on sterile coverslips of 12 mm diameter (Electron Microscopy Sciences, cat.no. 72230-01). Macrophages were stimulated with bacteria as described above and 15 ng LPS per 15,000 macrophages for 1 hr. After 1 h, RPMI-1640 media containing 10% FBS was supplemented with Gentamicin (Fisher Scientific, cat. no 12664735) to kill extracellular bacteria. A second antibiotic treatment was carried out for 2 h before macrophage lysis. At 4 hr post-infection (hpi), the macrophages were washed and fixed with 4% paraformaldehyde (Aurion, cat. no. 15710) and immunostained for Syntaxin 3 (Synaptic systems, cat no. 110033), LPS (Invitrogen, cat. no. PA1-73178), TNF-α (BioLegend, cat. no. 506302) or CD71 (clone b3/25). Secondary antibodies were anti-goat Alexa Fluor (AF)647 to detect LPS, anti-rabbit AF568 to detect syntaxin 3, and anti-mouse AF647 to detect TNF-α and CD71. Phalloidin AF488 (Invitrogen, cat. no. A12379) was used to stain F-actin and DAPI (Sigma Aldrich, cat. no. 32670) stained nucleic acids. Image acquisition and Z-stacks were done with a confocal laser scanning microscope (Zeiss, LSM 800) equipped with a 63× oil immersion objective lens.

### Transfections and TIRF microscopy

600,000 macrophages were cultured in a glass bottom culture dish (Greiner Bio One, cat. no. 627870) and transfected with Stx3-mCherry [13] or co-transfected with Stx3-mCherry (cloned as described (Verboogen *et al*., 2017)) and YFP-labeled TRPML1. TRPML1-YFP was a gift from Craig Montell (Addgene, #18826) (Venkatachalam *et al*., 2006). Briefly, 2 μg Stx3-mCherry alone or 1.7 μg each of Stx3-mCherry and TRPML1-YFPwas electroporated into macrophages by giving two pulses of 1,000 V with 40 ms width using a Neon Transfection system (Invitrogen).

Dried AF488 C5 maleimide (Invitrogen, A10254) was dissolved in ethanol to yield 50 μl of 2 mM reconstituted dye. An overnight culture of bacteria was washed and adjusted to 0.5 OD in 1 ml PBS. Labeling of bacteria occurred by incubating bacterial suspension with the reconstituted dye for 30 mins. After washing off the excess probe, the bacterial pellet was suspended in 2 ml PBS. For feeding macrophages, 400 μl labeled bacterial suspension was added to 600,000 transfected macrophages, yielding a ratio of approximately 10 macrophages per bacterium.

Total internal reflection Fluorescence (TIRF) live cell microscopy was carried out using a home-built TIRF microscope with an Olympus 60× UAPO NA 1.49 Oil objective and a Prime BSI Express sCMOS camera from Photometrics. Sequential excitation of the YFP and mCherry signals was used with alternating100 ms pulses of the 490 nm and 561 nm lasers. Excitation was split from the emission signals with a ZET405/488/561/647m dichroic mirror. The YFP and mCherry emissions were simultaneously imaged using an image splitter.

### Syntaxin 3 and TRPML1 knockdown

400,000 – 1,200,000 macrophages from 3 donors in 9 independent experiments were silenced using Syntaxin 3 siRNA: 5’ CCAAG CAGCU GACAC AGGAU GAUGA 3’ and 5’ UCAUC AUCCU GUGUC AGCUG CUUGG 3’. Briefly, 1.7 nmol duplex siRNA was electroporated into macrophages by giving two pulses of 1,000 V with 40 ms width using a Neon Transfection system (Invitrogen). 350,000 – 700,000 macrophages from 3donors in 3 independent experiments were electroporated with TRPML1 siRNA using a mixture. TRPML1 siRNA mixture was as follows: 5’ GGAGC AUUCG CUGCU GGUGA AUUGA 3’ and 5’ UCAAU UCACC AGCAG CGAAU GCUCC 3’; 5’ UCAUC CUGUU UGGGC UCAGU AAUCA 3’ and 5’ UGAUU ACUGA GCCCA AACAG GAUGA 3’; 5’ GAUCC GAUGG UGGUU ACUGA CUGCA 3’ and 5’ UGCAG UCAGU AACCA CCAUC GGAUC 3’. Briefly, 0.83 nmol siRNA duplex each from three siRNA pairs (i.e., a total of 2.5 nmol) was electroporated into macrophages as described above. In parallel, macrophages were electroporated with 1.7 nmol and 2.5 nmol medium GC duplex negative control siRNA (Fischer Scientific; cat. no 462001), respectively to be consistent with Syntaxin 3 siRNA and TRPML1 siRNA concentrations.

Following electroporation, the macrophages were cultured at a density of 15,000 macrophages per well for CFU assay in a 12-well plate at 500 μl per well and the remaining macrophages were seeded at ≥300,000 macrophages per well in a 6-well plate at 1 ml per well for parallel western blotting. This was done to check KD efficiency. Macrophages were stimulated with 15 ng LPS per 15,000 macrophages for 18 h at 37°C with 5% CO_2_. The macrophages seeded for western blotting were lysed after washing off the LPS. No bacteria were fed to these macrophages.

### ELISA on Syntaxin 3 KD

Syntaxin 3 silenced, and unsilenced macrophages were counted after electroporation to ensure seeding of 15,000 macrophages per well in a 12-well plate. The next day, macrophages were fed bacteria at a ratio of 50 macrophages per bacterium. After 1 h, RPMI-1640 media containing 10% FBS was supplemented with Gentamicin (Fisher Scientific, cat. no 12664735) to kill extracellular bacteria. A second antibiotic treatment was carried out for 2 h before macrophage lysis. At 4 hr post-infection (hpi), media was collected. IL-6 (Invitrogen; cat. no 88-7064) and TNF-α (Invitrogen; cat. no 88-7346) ELISAs were carried out according to the manufacturer’s instructions. Absorbance measurements at 450 and 570 nm were carried out using a microplate reader (BioTek, Synergy HTX).

### Intracellular colony forming unit (CFU) assay on Syntaxin 3 and TRPML1 silenced macrophages

Syntaxin 3 and TRPML1 silenced macrophages, and negative control siRNA transfected macrophages were counted after electroporation to ensure seeding of 15,000 macrophages per well in a 12-well plate. Unsilenced macrophages were also seeded at 15,000 macrophages per well. All macrophages were treated with 15 μl of 1 ng/μl LPS for 18 h at 37° C with 5% CO_2_. The next day, LPS was washed and the macrophages were fed bacteria at a ratio of 50 macrophages per bacterium. After 1 h, RPMI-1640 media containing 10% FBS was supplemented with Gentamicin (Fisher Scientific, cat. no 12664735) to kill extracellular bacteria. A second antibiotic treatment was carried out for 2 h before macrophage lysis. At 4 hr post-infection (hpi), the macrophages were washed and lysed using 0.1% Triton-X100. Lysates were cultured on LB agar plates at 37°C overnight.

In the case of Syntaxin 3 knock down, macrophages were fixed 4 h post-infection with 4% PFA for immunofluorescence microscopy. This was done by counting intracellular bacteria and comparing the result with the CFU assay.

### Intracellular CFU assays with TRPML agonist and calcium chelators

15,000 macrophages were seeded per well in 500 μl and activated with 15 μl of 1 ng/μl LPS for 18 h. The next day, LPS was washed off and the macrophages were treated with 2 μM MK6-83 (cat no. 5547, Tocris), 20 μM MLSA-1 (cat. no SML0627, Sigma), 20 μM SF22(cat. no.02120, Glixx laboratory Inc.) for 1 h. For calcium chelation, 5 μM BAPTA-AM (cat. no B6769, Invitrogen) and 3 mM EGTA (cat. no. 324626, Merck) was used for 1 h pre-treatments. An equal volume of DMSO was added as solvent control. All compounds were washed and the macrophages were cocultured with *E. coli* at 50 macrophages per bacterium. After 1 h, RPMI-1640 media containing 10% FBS was supplemented with Gentamicin to kill extracellular bacteria. Another antibiotic treatment for 2 h was carried out before macrophage lysis. At 4 hr post-infection, macrophages were washed and lysed using 0.1% Triton-X100. Lysates were cultured on LB agar plates at 37°C overnight.

### Western blotting

Control (unsilenced) and Syntaxin 3, TRPML1, and medium GC duplex negative control siRNA electroporated macrophages (≥300,000 per well in 1 ml) were stimulated with 15 ng LPS per 15,000 macrophages for 18 h at 37°C with 5% CO_2_. The next day, the LPS was washed off and macrophages were lysed by boiling at 95° C for 5 mins in a buffer containing 65.8 mM Tris HCl pH 6.8, 2.1 % SDS, 26.3 % (w/v) glycerol. Total proteins were quantified using the EZQ protein quantification kit (Invitrogen, cat. no: R33200). Equal amounts of proteins were resolved using 4-20% SDS-PAGE (Bio-Rad, cat. no: 456-1094), and blotted onto 0.2 μm PVDF membranes (Bio-Rad, cat no. 1620177). Blots were incubated overnight at 4°C with the following primary antibodies: Syntaxin 3 (Abcam, cat. no. 133750), TRPML1 (Novus Biologicals, cat. no. NBP1-92152), and GAPDH (Cell Signaling, cat. no. 2118). The secondary antibody was IR dye 680 nm conjugated donkey anti-rabbit (LI-COR Biosciences, cat. no. 926-32223). Imaging was done using channel 700 nm of Odyssey FC imager.

### Data analysis

All data is represented as mean ± SEM. Statistical analysis was carried out using Graph pad prism. P values ≤0.05 were considered significant. Protein band densitometry of western blots were done using Image Studio (LI-COR Biosciences). Confocal microscopy fixed images and Z-stacks were analyzed using Image J.

## Results

### Focal exocytosis of Syntaxin 3 to nascent phagosomes

Previously, it was shown in the murine dendritic cell line JAWSII and murine bone marrow-derived dendritic cells, that Syntaxin 3 translocates from intracellular compartments to the plasma membrane upon LPS stimulation, and this promotes the secretion of IL-6 and MIP-1α (Collins *et al*., 2015). We started by confirming these results in human peripheral blood monocyte-derived macrophages. In unstimulated macrophages, Syntaxin 3 was located largely in the perinuclear area (**Figure 1 and Supplementary Figure 1A**). However, upon stimulating the macrophages with LPS, we found that Syntaxin 3 translocated to the plasma membrane (**Figure 1A and Supplementary Figure 1A**), similar to the previous observations (Collins *et al*., 2015).

**Figure 1.**
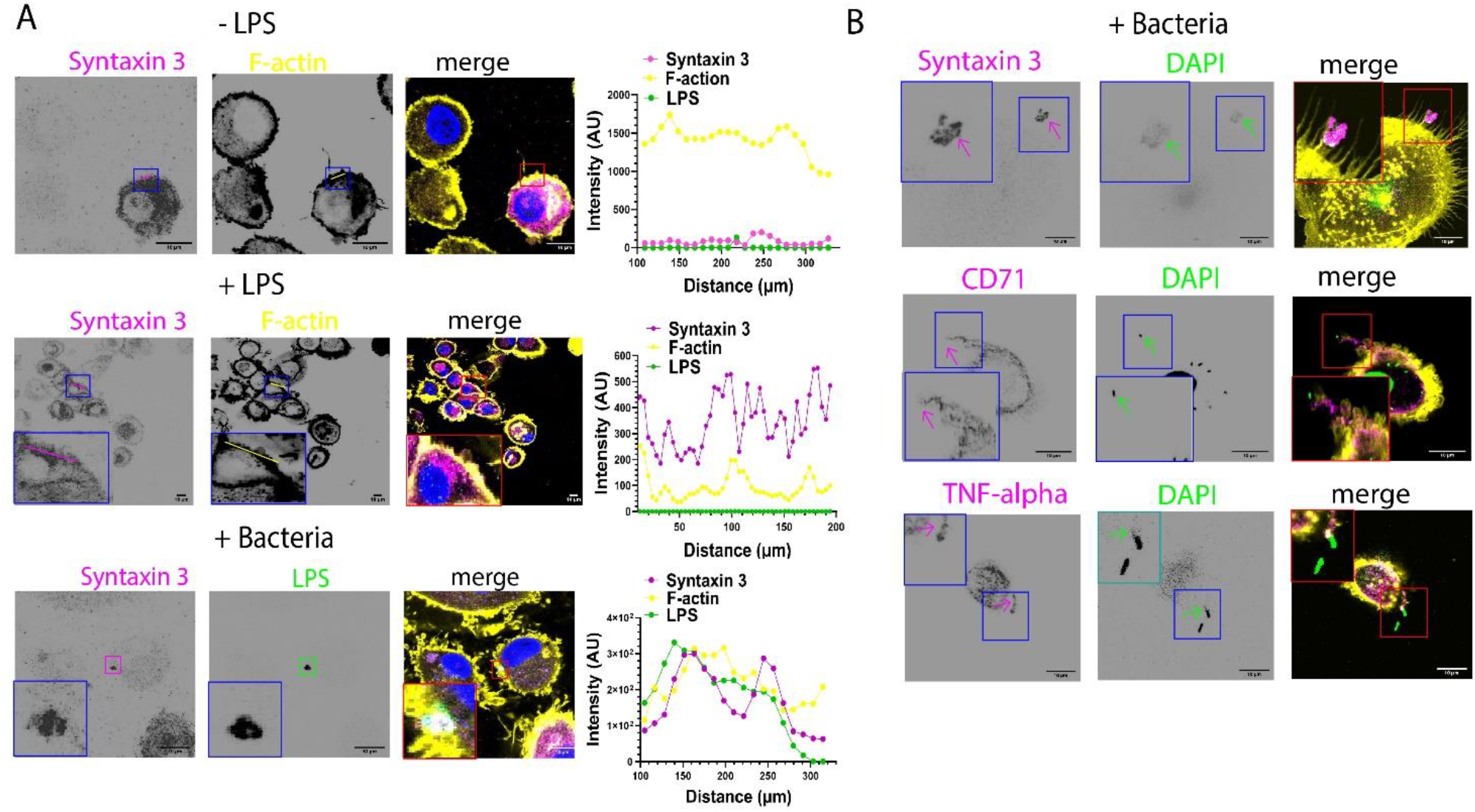
Polarized trafficking of syntaxin 3 to nascent phagosomes. (A) Immunofluorescence confocal microscopy images of unstimulated macrophages (upper panels), macrophages stimulated with 15 ng LPS (middle panels), and macrophages stimulated with *E. coli* (lower panels). Macrophages were stained for Syntaxin 3 (magenta), F-actin (yellow), and LPS (green). DAPI is in blue. (B) Immunofluorescence confocal microscopy images of macrophages stimulated with *E. coli* and stained for Syntaxin 3 (upper panes), CD71 (middle panels), and TNF-α (lower panels). **See also Supplementary Figures 1 -5**. Scale bars, 10 μm.

Gram-negative bacteria, such as *E. coli*, contain high amounts of LPS in their outer membrane. We, therefore, determined whether Syntaxin 3 would selectively translocate to nascent phagosomes of *E. coli* bacteria (i.e., the contact points with the bacteria). Indeed, upon stimulating macrophages with bacteria, Syntaxin 3 selectively translocated to the surface of the attached bacteria (**Figure 1B, Supplementary Figure 1C, and Supplementary Movies 1 – 3**). We found that 90% of macrophages showed polarization of Syntaxin 3 towards invading bacteria. Immunofluorescence microscopy on bacteria stimulated macrophages also suggest that SNAP23 might interact with syntaxin 3 at the plasma membrane since focal localization of SNAP23 is also observed at the nascent phagosomes of pseudopodia (**Supplementary Figure 1D**). As controls, we also determined focal exocytosis of recycling endosomes (Bajno *et al*., 2000), using the marker CD71, and TNF-α, which are also delivered to phagosomes in a polarized fashion (Manderson *et al*., 2007; Murray & Stow, 2014). 90% of macrophages showed polarization of CD71 and 70% showed polarization of TNF-α towards invading bacteria (**Figure 1B and Supplementary Figure 2A-C**).

### Syntaxin 3 is required for bacterial uptake

Next, we determined whether the focal exocytosis by Syntaxin 3 would be required for bacterial uptake. We silenced Syntaxin 3 using siRNA and determined the number of ingested bacteria per macrophage by microscopy. On average, we attained ∼60% knockdown efficiency (**Figure 2A**). While we observed less intracellular bacteria in Syntaxin 3 silenced macrophages for two out of three donors (**Figure 2B, left panel**), this difference was not significant, probably because bacteria are small and hence accurately counting bacteria in a single focal plane of a cell is technically challenging. Therefore, we also performed intracellular colony forming unit (CFU) assays. In these CFU assays, macrophages were first incubated with live bacteria for a short period. Subsequently, the extracellular (i.e. non-ingested) bacteria were killed with the membrane impermeable antibiotic gentamicin. Then, the macrophages were lysed and the intracellular bacteria were cultured on LB agar plates. This is a highly sensitive method for determining the uptake efficiency of *E. coli*, as individual intracellular bacteria form colonies on the agar plates. We observed that less bacteria were internalized by Syntaxin 3 silenced macrophages, relative to the non-targeted siRNA control (**Figure 2B, right panel**). Additionally, Syntaxin 3 silencing caused a significant decrease in bacteria-stimulated IL-6 secretion but not TNF-α secretion (**Figure 2C, D**), as reported previously (Collins *et al*., 2015).

**Figure 2.**
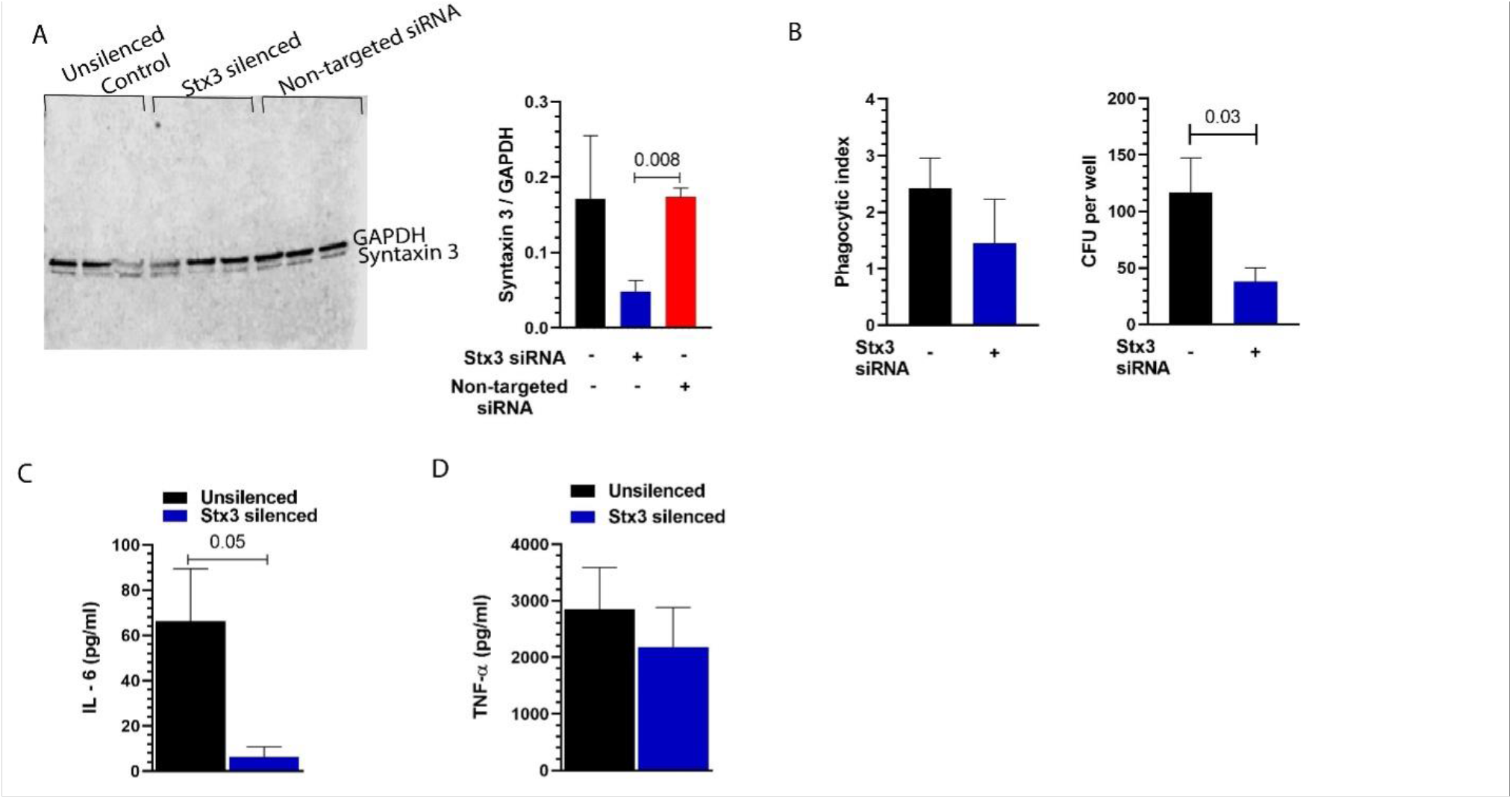
Syntaxin 3 silencing reduces bacterial uptake and IL-6 secretion by macrophages. (A) Representative western blot showing Syntaxin 3 and GAPDH bands in macrophages from 3 different donors, transfected with siRNA targeting Syntaxin 3 and non-targeting siRNA control. GAPDH: loading control. Right: Quantification of Syntaxin 3 levels. n = 3 - 8 donors. (B) Left: Bacterial uptake by Syntaxin 3 silenced macrophages as determined by immunofluorescence microscopy, n = 3 donors. Right: Intracellular colony forming unit assay (CFU) assay showing decreased bacterial uptake by Syntaxin 3 macrophages, n = 6 donors. (C) IL-6 secretion by Syntaxin 3 silenced macrophages stimulated with *E. coli* as determined by ELISA. n = 6 (D) Same as panel C, but now for TNF-α. Two-tailed unpaired t-test was used to determine statistical significance.

### Co-delivery of Syntaxin 3 and TRPML1 at pseudopodia of nascent phagosomes

To provide direct evidence that vesicles positive for Syntaxin 3 are focally delivered at nascent phagosomes, we used live cell TIRF microscopy of macrophages expressing Syntaxin 3 C-terminally labeled with the fluorescent protein mCherry. Indeed, we observed focal delivery of Syntaxin 3 vesicles at nascent phagosomes concomitant with an increase in the length of the pseudopodia (**Figure 3; Supplementary Movies 6, 7**). Such focal exocytosis at the forming phagosomes has been observed previously for TRPML1-positive lysosomes (Samie *et al*., 2013). Therefore, we next assessed whether Syntaxin 3-positive vesicles also contained TRPML1 using live cell TIRF microscopy with macrophages co-expressing Syntaxin 3-mCherry together with TRPML1 fused to the spectrally shifted fluorescent protein YFP (Venkatachalam *et al*., 2006). Indeed, we observed that Syntaxin 3-mCherry and TRPML1-YFP double positive vesicles were delivered to forming phagosomes (**Figure 4A, B, Supplementary Movies 8 – 11**).

**Figure 3.**
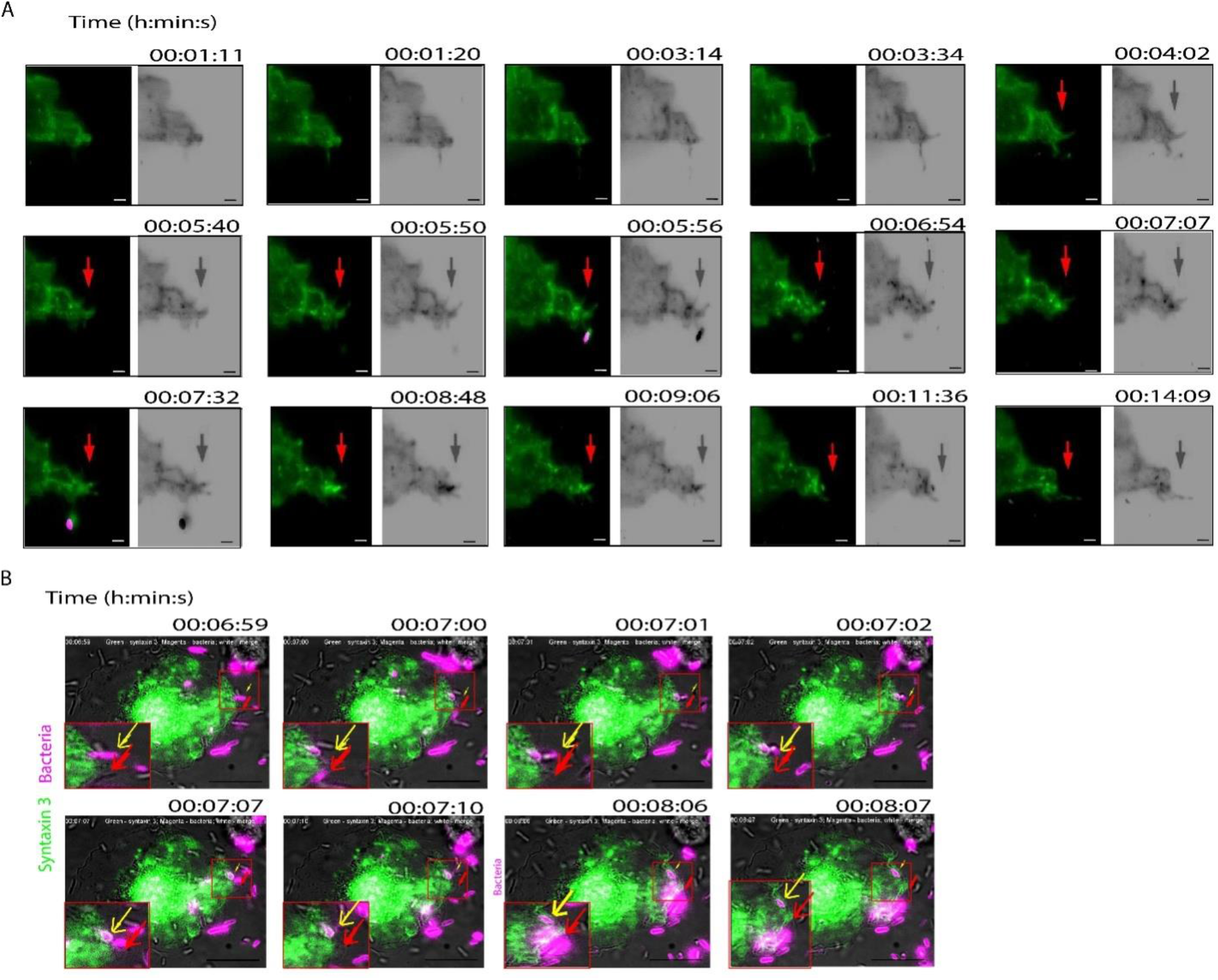
Focal exocytosis of Syntaxin 3 at nascent phagosomes. (A) Snapshots of time-lapse live cell TIRF microscopy of macrophages expressing Syntaxin 3-mCherry and pulsed with fluorescently labeled bacteria. Red arrows: delivery of Syntaxin 3-positive vesicles to the pseudopodia. Left: Merge with Syntaxin 3 in green and bacteria in magenta. Right: Syntaxin 3 and bacteria in grey scale. See also **Supplementary Movie 6**. (B) Same as panel A, but now only merge is shown. Yellow and red arrows show co-localization of Syntaxin 3 and phagocytosing bacteria at 7:00 and 8.06 mins, respectively. See also **Supplementary Movie 7**. Scale bars, 1 μm in panel A and 10 μm in panel B.

**Figure 4.**
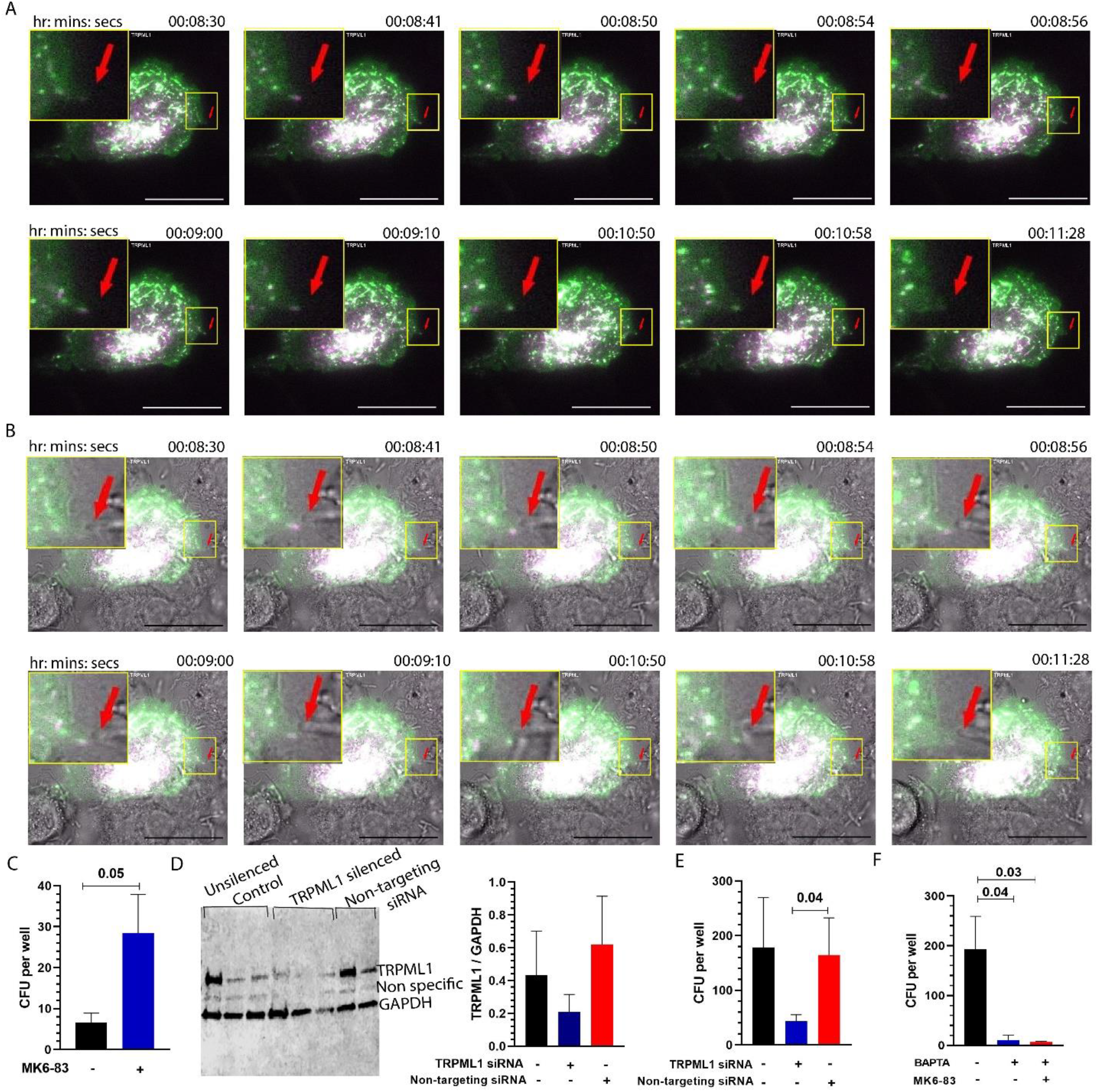
Delivery of Syntaxin 3 and TRPML1 to nascent phagosomes. (A) Snapshots of time-lapse live cell TIRF microscopy of macrophages expressing TRPML1-YFP (green) and Syntaxin 3-mCherry (magenta), and pulsed with unlabeled bacteria. Red arrow: focal delivery of TRPML1 and Syntaxin 3. (B) Same as panel A, but now also including the bright field channel. See also **Supplementary Movies 8 – 11**. (C) Intracellular colony forming unit (CFU) assay of macrophages treated with 2 μM of the TRPML1 agonist MK683. n = 5 donors. (D) Representative western blot showing TRPML1 and GAPDH bands in TRPML1 silenced macrophages. n= 3 donors. Note that expression levels of TRPML1 varies among different donors. GAPDH: loading control. Graph shows quantification of TRPML1 levels. (E) Intracellular CFU assay of macrophages with TRPML1 knock down. n = 7 donors. (F) CFU assay of macrophages after calcium chelation using 5 μM BAPTA. n = 3 donors. Two tailed unpaired t-test and ordinary one-way ANOVA is used to determine statistical significance. Scale bars,10 μm.

To confirm the role of TRPML1 in phagocytosis (Samie *et al*., 2013), we next pre-treated the macrophages with 2 μM of the TRPML1 agonist MK6-83 before incubating them with the bacteria and performing CFU assays. The number of intracellular bacteria significantly increased in MK6-83-treated macrophages (**Figure 4C)**, and a similar trend was observed in macrophages pre-treated with the other TRPML1 agonists MLSA-1 and SF22 **(Supplementary Figure 2D)**. To confirm the role of TRPML1 in bacterial uptake with an independent assay, we also silenced TRPML1 using siRNA and determined the number of ingested bacteria by the intracellular CFU assay. On average, we attained ∼50% knockdown efficiency (**Figure 4D**). The number of bacteria ingested by TRPML1 silenced macrophages significantly decreased relative to non-targeting siRNA control (**Figure 4E)**.

We also performed experiments with calcium chelation (i.e., the substrate of TRPML1), as we expected this would also inhibit phagocytosis of *E. coli* by the macrophages (Samie *et al*., 2013). The macrophages were pretreated with 5 μM of the intracellular chelator BAPTA-AM or 3 mM of the extracellular chelator EGTA alone and together with MK6-83 before feeding them bacteria (**Figure 4F and Supplementary Figure 2E**). Calcium chelation with BAPTA-AM completely blocked phagocytosis regardless of the presence of MK6-83. EGTA did not significantly affect bacterial uptake, but blocked the increase induced by MK6-83. These data indicate that focal exocytosis of vesicles carrying Syntaxin 3 and TRPML1 at the forming phagosomes promotes calcium transport, which in turn is required for phagocytosis.

## Discussion

In this study, we show that vesicles carrying Syntaxin 3 and TRPML1 undergo focal exocytosis at the plasma membrane site of phagocytosis in macrophages. Moreover, we show that both Syntaxin 3 and TRPML1 are required for the efficient uptake of *E. coli* bacteria. There are three, not mutually exclusive, ways how these data can be interpreted.

First, Syntaxin 3 might be involved in the delivery of TRPML1 at the plasma membrane, possibly by interactions with VAMP7 (Dingjan *et al*., 2018). In this case, the knockdown of Syntaxin 3 would interfere with the delivery of TRPML1 at the nascent phagosome and inhibit phagocytosis indirectly (i.e. via inhibition of TRPML1 activity).

Second, it could be that the functions of Syntaxin 3 and TRPML1 are not directly related to each other, but that both these proteins are required for phagocytosis independently. Syntaxin 3 has for example been described to interact with VAMP2 and VAMP3 (Dingjan *et al*., 2018), and these interactions might mediate the focal delivery of VAMP3-positive recycling endosomes (Bajno *et al*., 2000) and/or VAMP2-positive Golgi-derived vesicles (Vashi *et al*., 2017) independently of TRPML1 activity. In this case, both TRPML1 and Syntaxin 3 are thus co-delivered to the nascent phagosome by the same trafficking vesicles, but are not immediately functionally connected.

Third, TRPML1 might be required for the focal exocytosis of Syntaxin 3-positive lysosomes. We showed that the chelation of extracellular Ca^2+^ with EGTA had no significant effect on bacterial uptake, whereas chelation of intracellular Ca^2+^ with BAPTA-AM completely blocked uptake. These data suggest that influx of extracellular Ca^2+^ through plasma membrane-localized TRPML1 (and other Ca^2+^ channels) has no, or only a minor, contribution to phagocytosis, whereas the release of Ca^2+^ from intracellular compartments through TRPML1 located in intracellular compartments, is needed for efficient phagocytosis. Previously, TRPML1 has been shown to locate predominately at late endosomes and lysosomes (Pryor *et al*., 2006), and therefore, it seems likely that the Syntaxin 3 and TRPML1-positive compartments are of lysosomal nature. Lysosomes are a rich source of Ca^2+^ with an estimated intra-lysosomal concentration of ∼0.5 mM compared to a drastically lower cytosolic Ca^2+^ concentration of ∼100 nM (Xu & Ren, 2015). This would argue for a model where lysosome-located TRPML1 is activated at/near the site of phagocytosis, promoting the release of Ca^2+^ from the lysosomal lumen into the cytosol. This Ca^2+^ release in turn might trigger focal exocytosis of Syntaxin-3 carrying vesicles, for example via the Ca^2+^ sensor Synaptotagmin VII (Corrotte & Castro-Gomes, 2019).

Previously, it was shown that lysosome exocytosis occurs during the uptake of large particles (>3 μm), but not during the ingestion of small particles (Tardieux *et al*., 1992; Samie *et al*., 2013; Sun *et al*., 2020), presumably because the extra membrane supplied by the fusion of lysosomes with the plasma membrane provides additional membrane for the uptake of large particles. Indeed, lysosome exocytosis has been observed during ingestion of large trypanosomes (>10 μm) (Tardieux *et al*., 1992) and *Mycobacterium marinum* (1.5 – 4 μm) (Koo *et al*., 2008). Our data now suggest that lysosome exocytosis might also play a role in the uptake of ∼2 μm *E. coli*. This might relate to the other functions of focal lysosomal exocytosis at nascent phagosomes: (i) The repair of the injured plasma membrane that arises during pathogen invasion [18, 21], and (ii) the delivery of bactericidal enzymes, as lysosome contain ∼60 hydrolases which would weaken invading bacteria at the phagocytic cup (Xu & Ren, 2015).

We have previously proposed that vesicles or secretory lysosomes carrying calcium-independent phospholipase A_2_ beta (iPLA_2_-β) translocate to the forming phagosome to support membrane extension by promoting the local fusion of these vesicles with the plasma membrane (Dabral & van den Bogaart, 2021). Overexpression of PLA_2_ increases lysosome exocytosis and inhibition of TRPML1 drastically reduces PLA_2_-mediated lysosome exocytosis (Cui *et al*., 2021). This suggests that lysosome exocytosis is not only dependent on TRPML1 but also PLA_2_ activity (Cui *et al*., 2021). It would therefore be interesting to determine whether the Syntaxin 3 and TRPML1 carrying vesicles also contain iPLA_2_-β.

## Supporting information

Supplementary Movie 1. Surface co-localization of Syntaxin 3 and bacteria in an extented pseudopodium.

Supplementary Movie 2. Surface co-localization of Syntaxin 3 and bacteria in a retracted pseudopodium.

Supplementary Movie 3. Surface co-localization of Syntaxin 3 and bacteria.

Supplementary Movie 4. Intracellular co-localization of Syntaxin 3 and bacteria.

Supplementary Movie 5. Intracellular co-localization of syntaxin 3 and bacteria.

Supplementary Movie 6. Syntaxin 3 traffics to pseudopodia leading to its extension.

Supplementary Movie 7. Syntaxin 3 translocates to bacteria attachment site on the plasma membrane.

Supplementary Movie 8. Co-trafficking of Syntaxin 3 and TRPML1 to pseudopodia.

Supplementary Movie 9. Same as Supplementary Movie 8, but now with overlay of the bright field channel.

Supplementary Movie 10. Co-trafficking of Syntaxin 3 and TRPML1 to the nascent phagosomes.

Supplementary Movie 11. Same as Supplementary Movie 10, but now with overlay of the bright field channel.

## CRediT author statement

Deepti Dabral and Geert van Bogaart - Conceptualization; Deepti Dabral - Data curation; Deepti Dabral - Formal analysis; Geert van Bogaart - Funding acquisition; Deepti Dabral - Investigation; Deepti Dabral and Geert van Bogaart - Methodology; Deepti Dabral - Project administration; Geert van Bogaart - Resources; Deepti Dabral - Roles/Writing - original draft; Deepti Dabral and Geert van Bogaart - review & editing.

## Acknowledgments

GvdB has received funding from the European Research Council (ERC) under the European 8 Union’s Horizon 2020 research and innovation program (grant agreement No. 862137).

## Figures

**Supplementary Figure 1.**
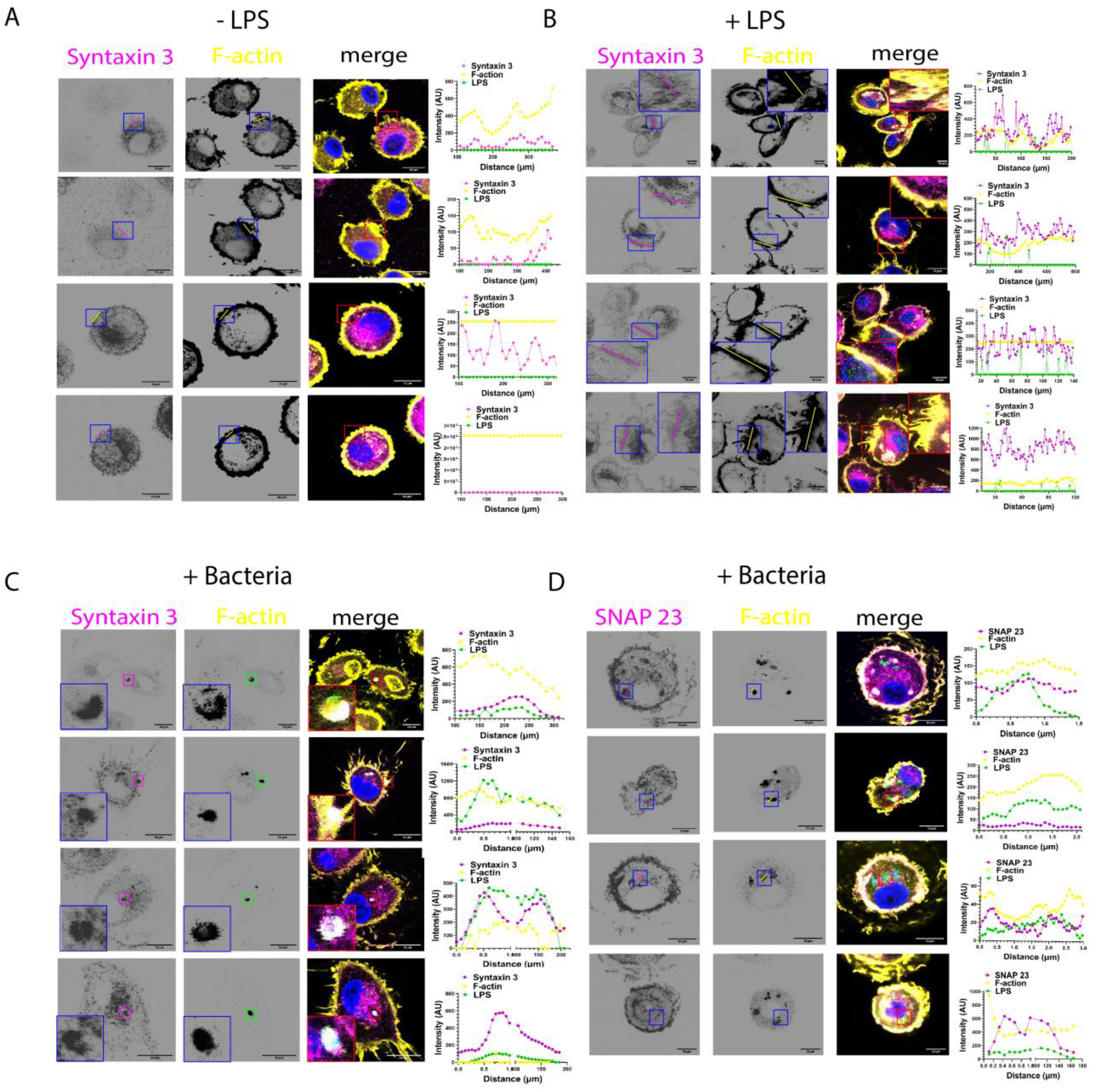
Panel of immunofluorescence confocal microscopy images showing location of Syntaxin 3 in (A) unstimulated macrophages, (B) macrophages stimulated with 15 ng LPS, (C) macrophages stimulated with *E. coli*. Magenta - Syntaxin 3. Yellow - F-actin. Blue - Nucleic acid. Green - LPS. Scale bars, 10 μm. (D) Panel of immunofluorescence confocal microscopy images showing location of SNAP23 stimulated with *E. coli*.

**Supplementary Figure 2.**
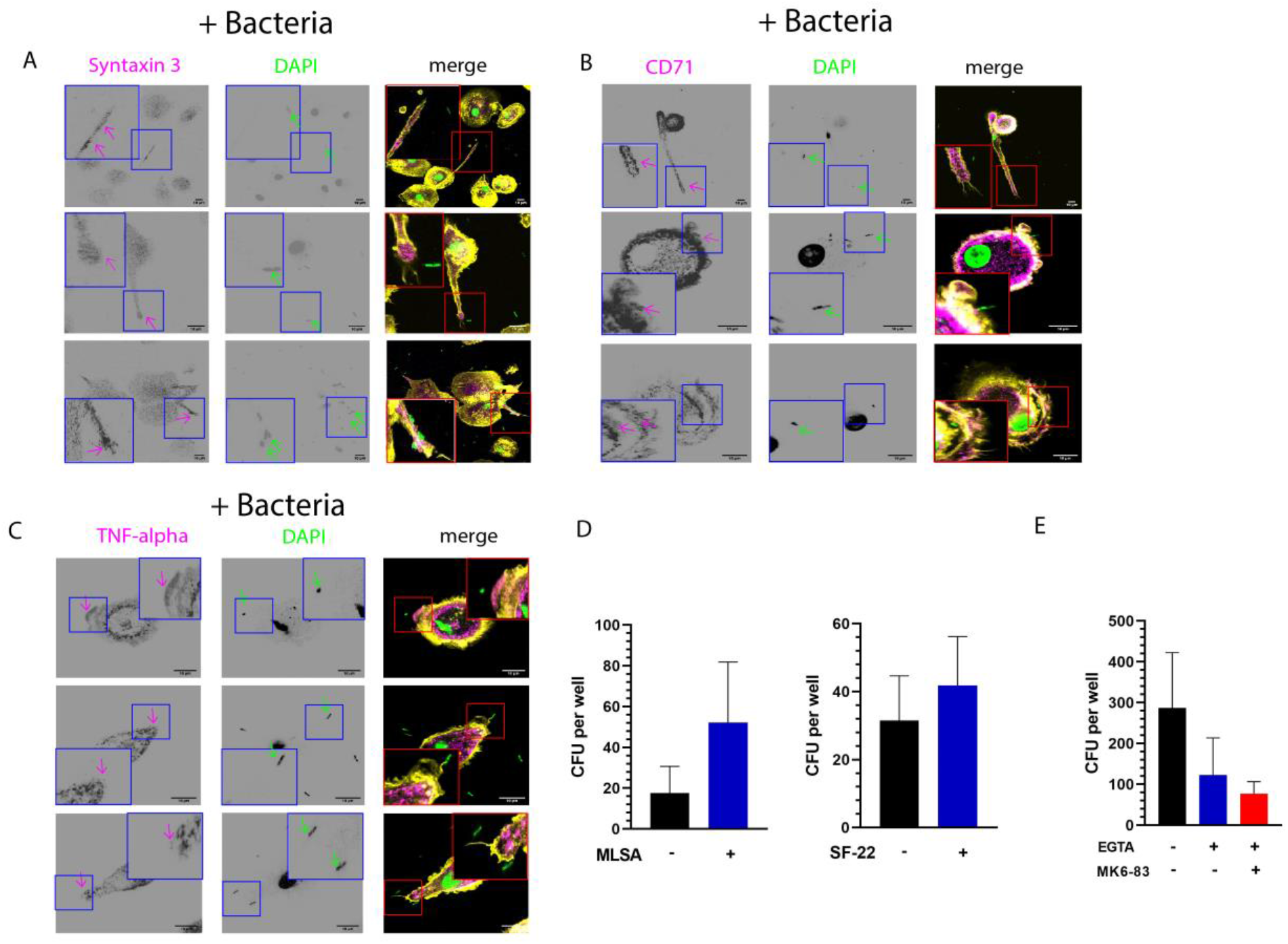
(A) Panel of immunofluorescence confocal microscopy images showing location of Syntaxin 3 in macrophages incubated with *E. coli*. Magenta: Syntaxin 3. Blue: DAPI. Yellow: F-actin. (B) Same as panel A, but now for CD71 instead of Syntaxin 3. (C) Same as panel A, but now for TNF-α instead of Syntaxin 3. (D) CFU assay showing bacterial uptake by macrophages treated with TRPML1 agonists 20 μM MLSA-1 and 20 μM SF22. n = 6 donors. (E) Chelation of calcium using 3mM EGTA reduce bacterial uptake. n = 3 donors. Scale bars, 10 μm.

**Supplementary Movie 1. Surface co-localization of Syntaxin 3 and bacteria in an extented pseudopodium**. Immunofluorescence confocal microscopy Z-stacks of a macrophage stimulated with *E. coli*. Macrophages were stained for Syntaxin 3 (magenta), F-actin (yellow), LPS (green) and DNA (blue).

**Supplementary Movie 2. Surface co-localization of Syntaxin 3 and bacteria in a retracted pseudopodium**. Immunofluorescence confocal microscopy Z-stacks of a macrophage stimulated with *E. coli*. Macrophages were stained for Syntaxin 3 (magenta), F-actin (yellow) and LPS (green).

**Supplementary Movie 3. Surface co-localization of Syntaxin 3 and bacteria**. Immunofluorescence confocal microscopy Z-stacks of a macrophage stimulated with *E. coli*. Macrophages were stained for Syntaxin 3 (magenta), F-actin (yellow) and LPS (green).

**Supplementary Movie 4. Intracellular co-localization of Syntaxin 3 and bacteria**. Immunofluorescence confocal microscopy Z-stacks of a macrophage stimulated with *E. coli*. Macrophages were stained for Syntaxin 3 (magenta), F-actin (yellow) and LPS (green).

**Supplementary Movie 5. Intracellular co-localization of syntaxin 3 and bacteria**. Immunofluorescence confocal microscopy Z-stacks of a macrophage stimulated with *E. coli*. Macrophages were stained for Syntaxin 3 (magenta), F-actin (yellow) and LPS (green).

**Supplementary Movie 6. Syntaxin 3 traffics to pseudopodia leading to its extension**. Time-lapse live cell TIRF microscopy of macrophages expressing Syntaxin 3-mCherry and pulsed with fluorescently labeled bacteria. Red arrows: delivery of Syntaxin 3 vesicles to the pseudopodia.

**Supplementary Movie 7. Syntaxin 3 translocates to bacteria attachment site on the plasma membrane**. Time-lapse live cell TIRF microscopy of macrophages expressing Syntaxin 3-mCherry and pulsed with fluorescently labeled bacteria. Yellow and red arrows show co-localization of Syntaxin 3 and phagocytosing bacteria at 7:00 and 8.06 mins, respectively.

**Supplementary Movie 8. Co-trafficking of Syntaxin 3 and TRPML1 to pseudopodia**. Time-lapse live cell TIRF microscopy of macrophages expressing TRPML1-YFP (green) and Syntaxin 3-mCherry (magenta), and pulse with unlabeled bacteria. Red arrow: focal delivery of TRPML1 and Syntaxin 3 to pseudopodia.

**Supplementary Movie 9**. Same as Supplementary Movie 8, but now with overlay of the bright field channel.

**Supplementary Movie 10. Co-trafficking of Syntaxin 3 and TRPML1 to the nascent phagosomes**. Time-lapse live cell TIRF microscopy of macrophages expressing TRPML1-YFP (green) and Syntaxin 3-mCherry (magenta), and pulse with unlabeled bacteria. Red arrow: focal delivery of TRPML1 and Syntaxin 3 at a nascent phagosome.

**Supplementary Movie 11**. Same as Supplementary Movie 10, but now with overlay of the bright field channel.

